# Endophytic Virome

**DOI:** 10.1101/602144

**Authors:** Saurav Das, Madhumita Barooah, Nagendra Thakur

## Abstract

Endophytic microorganisms are well established for their mutualistic relationship and plant growth promotion through production of different metabolites. Bacteria and fungi are the major group of endophytes which were extensively studied. Virus are badly named for centuries and their symbiotic relationship was vague. Recent development of omics tools especially next generation sequencing has provided a new perspective towards the mutualistic viral relationship. Endogenous virus which has been much studied in animal and are less understood in plants. In this study, we described the endophytic viral population of tea plant root. Viral population (9%) were significantly less while compared to bacterial population (90%). Viral population of tea endophytes were mostly dominated by endogenous pararetroviral sequences (EPRV) derived from Caulimoviridae and Geminiviridae. Subclassification of Caulimoviridae showed the dominance of *Badnavirus* (42%), *Caulimovirus* (29%), *Soymovirus* (3%), *Tungrovirus* (3%), while Geminviridae was only represented by genus Bagmovirus. Interestingly, the endophytic virome sequence from root also showed the presence of phage virus from order Caudovirales. Identified sequence from Caudovirales were *Myoviridae and Siphoviridae*. Sequence comparison with viral population of soil and root showed the possibility of horizontal transfer of Caudovirales from soil to root environment. This study will expand the knowledge on endogenous viruses especially for tea plant. This study will also help us to understand the symbiotic integration of viral particle with plant which could be used in broader sense to tackle different agronomic problems.

**Significance Statement:** Virus were badly named for centuries and mostly known for their disease-causing abilities. But recent development of omics tools has focused another facet which is symbiotic. This paper discusses about viral community identified from shotgun sequence of tea root samples which are endogenous in origin. Interestingly, we also identified sequences of phage virus from Caudovirales family which possibly have transmitted from soil. Here we also compared the soil virome community with tea virome to establish the hypothesis. This research will broaden the current knowledge on symbiotic relationship of virus and plant.

## 1. Introduction

Plant health in the natural environment depends on a plethora of interaction between macro and microorganisms. Endophytes are the diverse microbial community residing within plant tissues in a commensal manner. Scientific studies have proven the abilities of endophytes to benefit host plants in adaptation to diverse agroclimatic conditions including wide biotic and abiotic stresses. Endophytes are also known for plant growth promotion by production of metabolites that improves the nutrient acquisition(1). The root endophytes are of great interest for their close proximate association with soil. There are reports of diverse bacterial and fungal association with root as root endophytic community (2–4). However, virus which were mostly considered and studied for their disease-causing ability in plant, animals, and human had remained as a bad name till recently. Recent advancement in omics tools has abled to entice attention towards symbiotic viral host-interactions. Studies highlighted the co-evolution of virus with their hosts and genomic association in an asymptomatic manner (5, 6). The big unknown of viral diversity and symbiotic association needs further scientific attention.

Symbiotic relationships are ubiquitous straddling all domains of life. It was 19^th^ century when term “symbiotic” came into first use to describe lichen or association between two dissimilar entities. Symbiotic relationship falls on a continuum between mutualistic and antagonistic, where environment affects its conditional mutualism(7). Viruses are abundant and diverse biological entities on earth. Recent studies discovered their abundance in desert, ocean, soil, mammalian gut, and plant ecosystems. Ecological surveys highlighted the wide misconception associated with viral interaction(5, 6). Viral interaction may not be symptomatic all the time and it could be mutualistic also. Interaction among viruses and their respective hosts are dynamic and conditional. Viral symbiosis can lead to symbiogenesis, where the evolution of new species occurs through genomic fusion of two entities. Coevolution and symbiogenesis can be supported by the overwhelming abundance of viral sequences in extant genomes (8, 9). Coevolution of virus and host is also a major player in the diversification of life on earth. In plant virus biodiversity studies, viruses were found in thousands of plants without any symptomatic presence (10, 11). Virus ameliorates the impacts of biotic and abiotic stresses. Virus improves cold and temperature tolerance along with disease-suppressive efficiency of plants. In Yellowstone National Park, plants adapted to geothermal areas with efficiency to tolerate high temperature was found to be associated with a fungus which in turn infected with virus (12). In white clover, a persistent plant virus, *White clover cryptic virus* can prevent the formation of nitrogen-fixing nodule when there is an adequate amount of nitrogen in the soils as a part of energy conservation (6).

Lack of universal coding sequence such as ribosomal RNA gene found in all cellular life makes it challenging to study the diversity of virus. Metagenomic studies with shotgun sequencing have provided a tool to understand the huge wealth of viral information from different environmental samples (10, 13). The present study describes the viral sequences identified from shotgun metagenomic sequences of root endophytes of tea plant. Endophytic viral sequences were also compared with the native soil viral community to observe the selective association or any horizontal transmission from soil to root of the plant. A comprehensive discussion was provided to hypothesize the possible functional role for such symbiotic association. This study is important to uncover the broader prospect of endogenous viral association. Understanding the viral association with plant and their diversity will help us to address disease emergence in plant and to identify the beneficial integration, which could be used to tackle different agricultural issues.

## 2. Methods

Tea root samples were collected from non-infected tea plants from tea garden of Assam Agricultural University, Jorhat district, Assam, India (latitude: 26.757874 & longitude: – 94.209824). Soil sample was collected from the same tea garden from a depth of 0 – 6 inches. Collected root samples were thoroughly washed with water to remove the soil and other debris. Then the root samples were surface sterilized with frequent washing of sodium hypochlorite and ethanol. The surface sterilized root samples were used for the metagenomic DNA extraction. Total DNA from root sample was extracted using the Qiagen DNeasy plant mini kit (Qiagen, USA). Soil metagenome was extracted using MoBio Power soil DNA kit (MoBio, USA). Shotgun metagenomic sequencing was done using Illumina MiSeq platform with 2 x 300 libraries. The library was prepared with TrueSeq DNA Library Preparation Kit. The unprocessed sequence reads were uploaded and analyzed using MG-Rast server. The taxonomic annotation was done using the RefSeq library with default parameters (e-value = 5, % - identity = 60, and minimum abundance of 1 with representative hits). The virome orf’s were identified with source filtering option by removing eukaryotic, bacterial and archaeal sequences in MG-Rast server. The visualization and analysis were done using excel chart, krona charts and R statistics. Heatmap was constructed with R statistics using Bray Curtis dissimilarity method to show the differences in diversity between soil and endophytic virome sequences. Venn diagram was prepared with the software venny 2.0 to map common population in soil and root endophytic virome. The data files are available through MG-Rast Server: root endophyte: https://bit.ly/2I4Cy36; soil sample: https://bit.ly/2WRgV9Q)

## 3. Results and Discussion

Tea Root endophytes were dominated with bacterial orf’s (90%), followed by virus (9%) and archaea (1%) (**Figure 1a**). Identified virome sequences were mostly dominated with *Incertae sedis* phylum (100%). *Incertae sedis* is used in taxonomy when the broader relationship of the group is unknown or undefined. Division based on order showed two groups, one is unclassified (94%) and another is *Caudovirales* (6%) (**Figure 1b**). *Caudovirales* are group-I viruses with double-stranded DNA (dsDNA) with an icosahedral head attached to a tail by connector protein. Unclassified sequences were mostly derived from *Caulimoviridae* and *Geminiviridae* family (**Figure 1c**). *Caulimoviridiae* are double-stranded DNA virus. They are extensively reported for their endogenous pararetroviral sequences (EPRVs). They have also been reported for their natural integration into the host genome(14, 15)’(16). This natural integration with the host genome also suggests their co-evolution with the plant-virus pathosystem(16). Earlier, six classes of pararetrovirus have been identified, with genome size ranging from 7.5 – 9.3 kb viz. icosahedral *Caulimovirus, Soymovirus, Cavemovirus, Petuvirus,* and bacilliform *Badnavirus, Tungrovirus* (15). However, with the rapid development of sequencing technology, different types of EPRV have been discovered in various plant genomes. Recently, bacilliform Ordenoviruses and the icosahedral Solendovirus has been included in the group. A newly discovered virus, *Rose yellow vein virus* (RYVV) has also been included in the group of EPRV (17). Origin of EPRV often regarded as an accidental origin, but their presence has major consequence for the host plant. Their presence may contribute to the genome size modification, change of methylation pattern of the host genome, and they can also act as genomic reorganizers by induction of chromosomal rearrangement(7, 15, 17). EPRV has been reported from many plants including tobacco, petunia, tomato, rice, banana etc. It is assumed that EPRV can serve as homology-dependent gene silencing of the non-integrated counterpart viruses like *petunia vein clearing virus* (PVCV, genus *Petuvirus*) in petunia plant and *Tobacco vein clearing virus* (TVCV; genus *Cavemovirus*) in tomato plants (18). The same mechanism is also recorded from the wild banana plant (diploid *Musa balbisiana*) having *Banana streak virus* (BSV; genus *Badnavirus*) shows resistant to BSV (19). Endogenous integration of Geminiviral DNA has also been reported form the genome of *Nicotiana* tabacum and related *Nicotiana* species including *N. tomentosiformis, N. tomentosa* and *N. kawakamii*. This integration suggests their unique common ancestral event where their genome has acquired the Geminiviral DNA(20).

**Figure 1:**
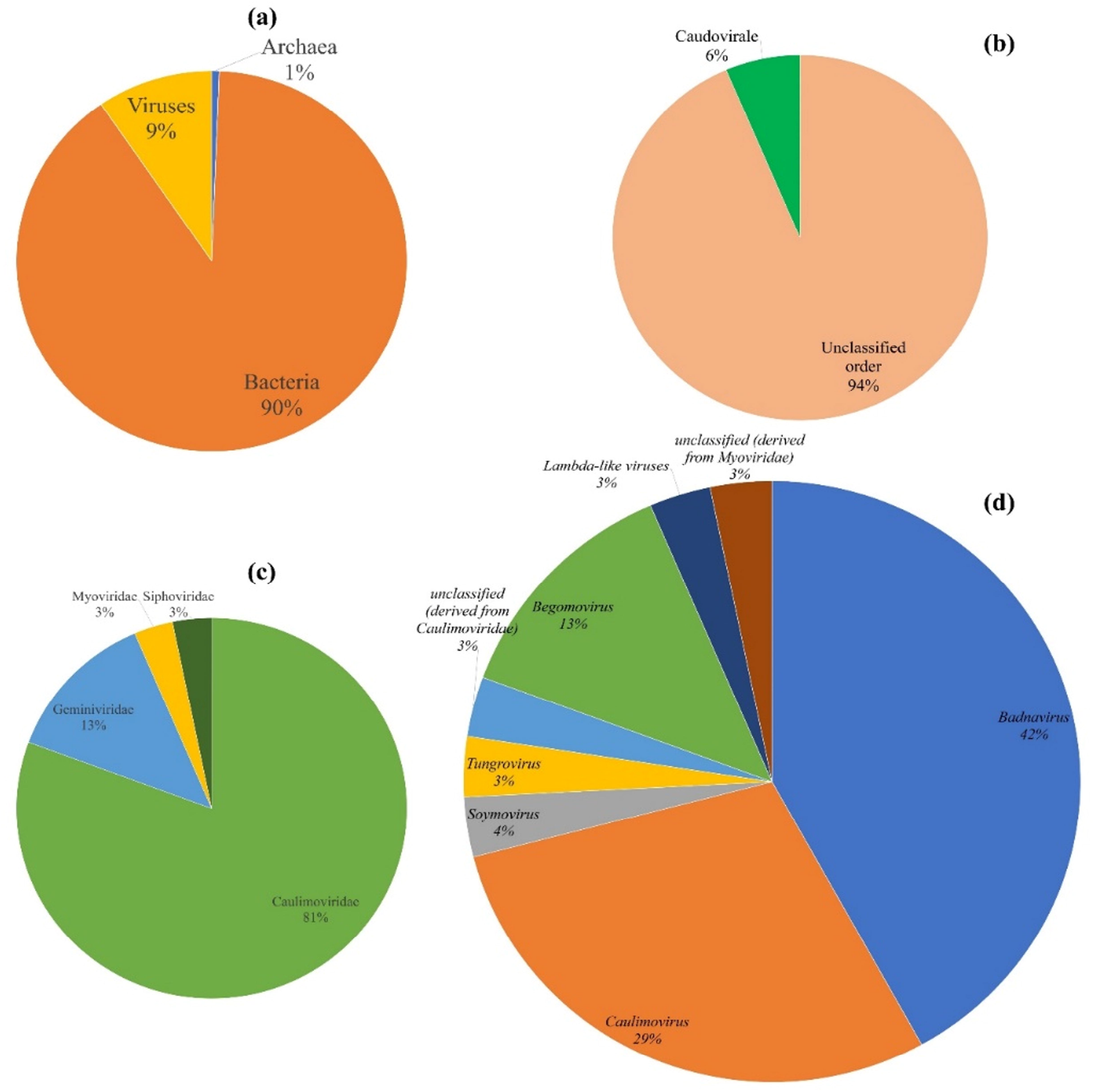
Tea endophytic Virome. (a) virus represent 9% of the total endophytic population between bacteria (dominant; 90%) and archaea (1%), (b) subclassification to order level showed two group Caudovirales and unknown order (c) Family wise classification showed a dominance of Caulimoviridae (81%) followed by Geminiviridae (13%) and Myoviridae (3%), Siphoviridae (3%).

Subclassification of Caulimoviridae to genus level showed sequence similarity with *Badnavirus* (42%), *Caulimovirus* (29%), *Soymovirus* (3%), *Tungrovirus* (3%), and unclassified sequences (3%) of *Caulomoviridae*. While *Geminiviridae* were mostly *Begomovirus* (13%) (**Figure 1d**). Most of the identified viral sequences were reported as pararetrovirus except *Begomovirus*. But this report could suggest their association as an endogenous viral particle as endogenous association of Geminiviridae is well known fact. *Badnavirus* is a plant associated bacilliform DNA virus of *Caulimoviridae* family. They have emerged as a serious pathogen to several horticultural crops like banana, black pepper, cocoa, citrus, sugarcane, taro and yam (21). *Badnavirus* are known for plant diseases like leaf chlorosis, root necrosis, red vein banding in young leaves, small mottled pods, and stem/root swelling followed by die-back (21, 22). Endogenous association of *Badnavirus* is reported by several researchers(19, 23). However, endogenous recombination with host genome may not cause any infection in host and can provide defense against non-integrative counterparts (21). There is only one report of *Badnavirus* form Tea plant that is also from metagenomic sequence of leaves and shoots samples by Hao et al., (2018) from China (24). *Caulimovirus* are plant pathogens and known for a disease like vein clearing and banding mosaic (25). *Soymovirus* is a dsDNA virus with worldwide distribution and few of the known species from the genus are *Soybean chlorotic mottle virus, Blueberry red ringspot virus, Cestrum yellow leaf curling virus,* and *Peanut chlorotic streak virus*. *Tungovirus* are a bacilliform virus of *Caulimoviridae*. They are mostly reported from monocots and *Poaceae* family. *Tungro bacilliform virus,* the sole member of the genus identified till now is known for rice tungro disease. Eastern coastal rice-growing region has a number of reports on tungro disease(26). They are also reported as pararetrovirus from rice. The genus *Begomovirus* in the family Geminiviridae, has wide host range infecting the dicot plants. Worldwide they are known for significant economic damage to may important crops like tomatoes, beans, squash, cassava and cotton. They are not reported from tea plant yet. The sequence of *Caudovirales* got subdivided into *Siphoviridae* and *Myoviridae* family. *Siphoviridae* are dsDNA virus and bacteria and archaea serve as their natural host. Genus wise classification showed unclassified sequence of Myoviridae and *Lambda-like virus* from *Siphoviridae* family.

Next question we wanted to address, is there any link between the viral population of soil (**Figure 2a**) from the tea garden and endophytic virome (**Figure 2b**)? But interestingly, there is no associated link between the soil viral population and tea endophytic virome except the sequences identified from order *Caudovirales (Siphoviradae* and *Myoviradae*) (**Figure 2c**). This also defines the endogenous origin of identified viral sequences. Soil viral sequences were mostly dominated with Unclassified genus of *Siphoviradae* and *Myoviridae, T4- like virus, P2- like virus*, *lamda like virus*, *Chlorovirus*, *Chlamydiamicro virus* etc. The presence of *Caudovirales* DNA sequences either as episomal/integrated possibly indicate the endophytic nature of these viruses. The transmission of *Caudovirales* from soil to endophytic root may have been mediated by soil-inhabiting bacteria or archaea. However, the compatibility and stability of viral sequences within the plant tissues for long term persistence needs to be verified. As plant didn’t serve as natural host for Siphoviradae and Myoviradae virus, they didn’t cause symptomatic infection in the tea plant.

**Figure 2:**
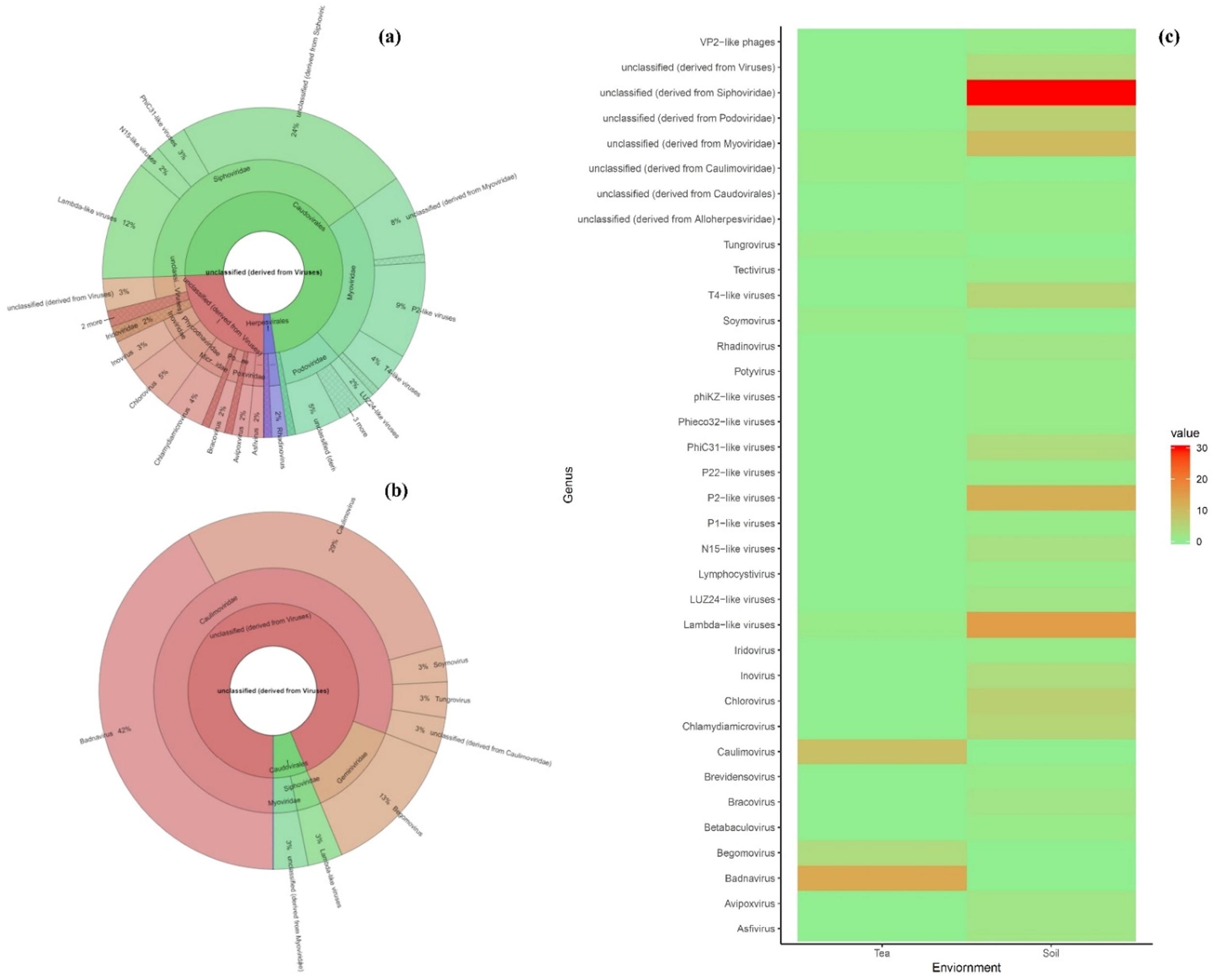
Diversity and divergence of viral population between soil and root endophytes. (a) Soil viral population (b) Tea endophyte: root virome (c) Heatmap: correlation between the soil and tea root endophytic viral population.

The mapping between the population of soil and root endophytic virome showed 94.4% of the identified endophytic genome were endogenous in origin and interestingly all are dsDNA virus (**Figure 3**). Except the *Caudovirales* which has originated from soil, possibly all the *Caulimoviradae* and *Geminiviridae* have coevolved with the tea plant as endogenous entities or as EPRV. Though there are ample studies on EPRVs, the contribution of EPRVs to plant genome and beneficiary effect is still poorly understood. The most accepted hypothesis is defensive mechanisms against infection or improves adaptability in stressed environment. It has been theorized that genome plasticity has allowed plants to acquire new biochemical process to adapt in competitive and variable climatic condition. The tea plant grows better in acidic soil where pH is 5.0 – 6.2. Adaptability in acidic soil may have been conferred by viral association as observed in the plants nearby geothermal soils of Yellowstone National Park. Viral association with the plants at geothermal soils helps in ameliorating the effect of thermal abiotic stress (> 50 °C). The grass plant showing resistance at Yellowstone National Park were colonized by a fungal endophyte which in turn infected with a virus. This three-way mechanisms between fungi, virus, and plant help them to survive in the geothermal soil. Viruses were also found to confer drought and cold tolerance in plants. Virus may also regulate the biosynthetic pathway of flavonoids and antioxidants in Tea plant. Even in natural integration, infection of clover plant with asymptomatic persistent *White clover cryptic virus,* can suppress the formation of nitrogen-fixing nodules when adequate nitrogen is present in the soil. Plant virus can also impact biotic stress factor, like white clover infected with *White clover mosaic virus* makes the plant less attractive fungus gnats (5). Infection of wild gourds with *Zucchini yellow mosaic virus* reduces the production of volatile compounds that attract the beetles to the plants (27). Viruses are a continuum between the mutualism and antagonism, while its active role always depends on the environment.

**Figure 3:**
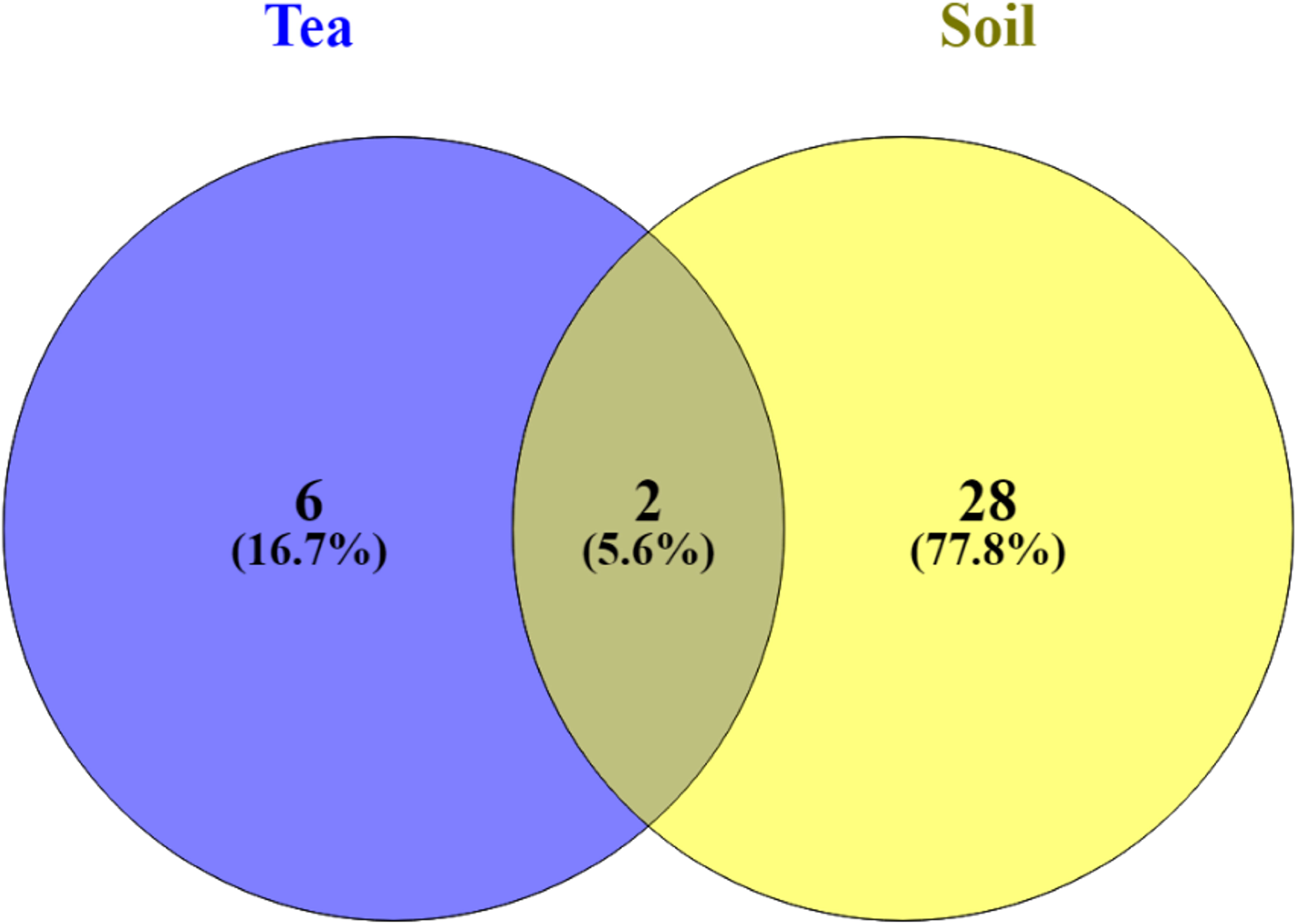
Association between tea root endophytic virome and soil virome sequence. Soil has highest number of viral sequences (77.8%) compared to root endophyte (16.7%) out of total population. However, 94.4% (considering 100% for whole endophytic population – 5.6% (2 common genus between soil and endophyte)) sequence of the endophytic virome were unique root endophyte and they are endogenous in origin.

Metagenomic era has greatly expanded the knowledge of plant virus. Plant virus serves as an important factor in all aspects of plant interaction with environment. Like fungi, bacteria, insect, virus is also an important biotic component of the plant’s environment. Understanding of viruses in ecological perspectives will enlighten the intertwined relationships of all life, including viruses. This study provides taxonomic distribution of viral endophytes in tea roots and discusses their possible role in different metabolic or defensive mechanisms. Tea root endophytes were mostly dominated with *Caulimoviridae, Caudovirdiae* and *Geminivridae* family. *Caulimoviridae* and *Gemivirdae* are endogenous in origin, while *Caudoviradae* have transmitted from soil. This report will expand the current knowledge on endogenous viruses especially for tea plant. There will be much more mutualistic virus which needs further study to understand their role in evolutionary perspectives.

## Acknowledgement

Authors would like to acknowledge Dr. Dipak Santra, Associate Professor, Department of Agronomy and Horticulture, University of Nebraska – Lincoln, USA for his suggestion and advice during manuscript preperation. Authors also extend their gratitude towards the faculty members of Department of Biotechnology, Assam Agricultural Univeristy for their continuous support with chemicals and instrument for carring out the experiments.

## Authors contribution

SD and MB designed the study; SD & NT prepared the draft; SD, NT & MB reviewed and finalized the draft.

